# Optogenetic stimulation of VIPergic SCN neurons induces photoperiodic changes in the mammalian circadian clock

**DOI:** 10.1101/2021.01.04.425287

**Authors:** Michael C. Tackenberg, Jacob J. Hughey, Douglas G. McMahon

## Abstract

Circadian clocks play key roles in how organisms respond to and even anticipate seasonal change in day length, or photoperiod. In mammals, photoperiod is encoded by the central circadian pacemaker in the brain, the suprachiasmatic nucleus (SCN). The subpopulation of SCN neurons that secrete the neuropeptide VIP mediate the transmission of light information within the SCN neural network, suggesting a role for these neurons in circadian plasticity in response to light information that has yet to be directly tested. Here, we used *in vivo* optogenetic stimulation of VIPergic SCN neurons followed by *ex vivo* PERIOD 2::LUCIFERASE (PER2::LUC) bioluminescent imaging to test whether activation of this SCN neuron sub-population can induce SCN network changes that are hallmarks of photoperiodic encoding. We found that optogenetic stimulation designed to mimic a long photoperiod indeed altered subsequent SCN entrained phase, increased the phase dispersal of PER2 rhythms within the SCN network, and shortened SCN free-running period – similar to the effects of a true extension of photoperiod. Optogenetic stimulation also induced analogous changes on related aspects of locomotor behavior *in vivo*. Thus, selective activation of VIPergic SCN neurons induces photoperiodic network plasticity in the SCN which underpins photoperiodic entrainment of behavior.

## Results

In mammals, circadian responses to seasonal changes in light cycles, or photoperiods, are encoded by the mammalian master clock, the suprachiasmatic nucleus (SCN) of the hypothalamus [1]. How the SCN initiates network changes due to photoperiod and maintains those responses, though, is not fully understood. The subpopulation of SCN neurons that express vasoactive intestinal polypeptide (VIP) has several functions that suggest a role in photoperiodic responses. VIPergic neurons relay photic information from the retina, even when their individual cellular clocks are deficient [2], and their activity is required for clock resetting from light exposure [3]. VIP itself is necessary for the induction of locomotor behavior duration after-effects [4]. Our lab has previously used SCN-targeted optogenetics to manipulate circadian behavior *in vivo*, and others have expanded this work to specifically target optogenetic stimulation to the VIPergic SCN [5]. Considering that long photoperiods extend the high-firing phase of the daily electrical rhythm of the SCN [6], we hypothesized that light-driven activation of VIP neurons may be a key step in photoperiodic encoding and SCN plasticity. Here we tested that hypothesis by stimulating VIPergic SCN neurons *in vivo* for an extended duration in mice entrained to a short photoperiod and then assaying effects on the SCN clockworks *ex vivo*.

We generated mice expressing both the PERIOD 2::LUCIFERASE (PER2::LUC) reporter and VIP-driven channelrhodopsin (VIP-ChR2), as well as non-ChR2 PER2::LUC control mice (ChR2-Neg), then implanted each mouse with a fiber optic targeting the SCN. We exposed these mice to 8 h of light, 8 h of fiber optic stimulation in the dark, and 8 h of unstimulated dark each day for 7 days. We then extracted and cultured 300-μm SCN slices and measured PER2::LUC expression for 6 days using an ICCD camera.

Using the peak times of the PER2::LUC rhythm for each of several thousand ~10 × 10 μm regions-of-interest (ROIs) from each SCN, we constructed a relative phase map for each slice at each peak (Figure 1A, Figure S1-2). These maps indicated that phase variation increased in both groups over time and that medial regions of the SCN became phase advanced relative to lateral regions. To quantify the changes in phase variation, we combined peak times for all ROIs from a given group (Figure 1B) and calculated the median absolute deviation for each slice for every cycle (Figure 1C). VIP-ChR2 slices had a significantly wider phase distribution compared to ChR2-Neg slices (*p* = 0.010, Two-Way ANOVA main effect of group), with a significant increase over time for both groups combined (*p* < 0.0001, Two-Way ANOVA main effect of peak). The change in SCN phase distribution in response to VIPergic SCN neuron activation therefore matches that previously observed following exposure to a long photoperiod compared to short [7].

**Figure 1.**
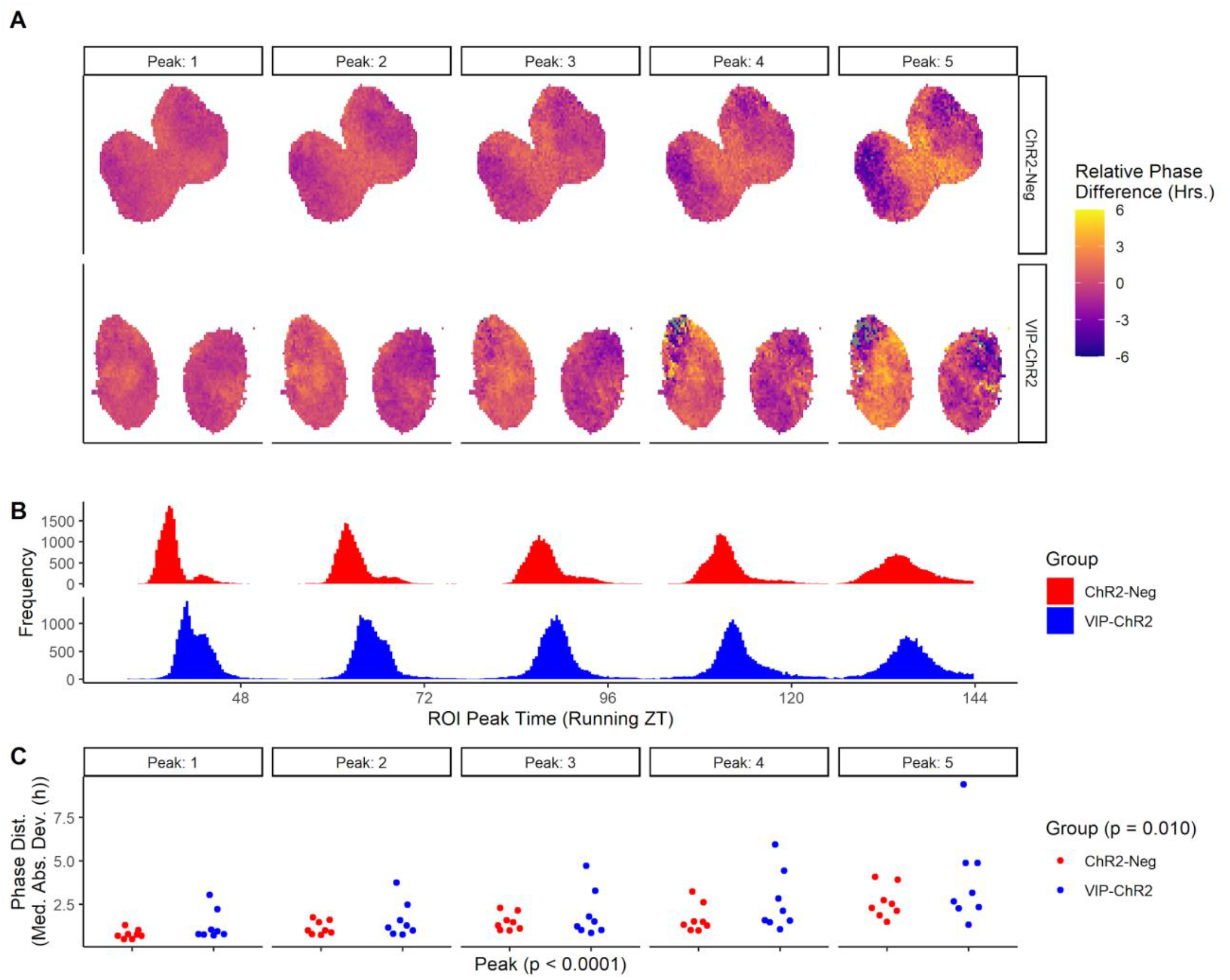
Optogenetic stimulation of VIPergic neurons influences *ex vivo* circadian phase distribution. A, representative heat maps showing the relative phase of the PER2::LUC rhythm for each ROI, for each peak (see also Figure S2). B, frequency distribution of the ROI peak time of each group (see also Figure S1). C, the median absolute deviation of the peak time for each peak in each slice. Peak and Group *p* values indicated are the main effects of an ordinary two-way ANOVA.

We next measured the free-running period of the overall PER2::LUC rhythm in the SCN slices from each group using the Lomb-Scargle periodogram (LSP, Figure 2A) [8]. PER2::LUC free-running period was significantly shorter (*p* = 0.001, Kruskal-Wallis test) in slices from VIP-ChR2 mice (median period 23.68, IQR 0.38 h) than in slices from ChR2-Neg mice (median period 24.42, IQR 0.23 h). The effect of VIPergic SCN neuron stimulation therefore mimics the effect of long photoperiod exposure *ex vivo* [7].

**Figure 2.**
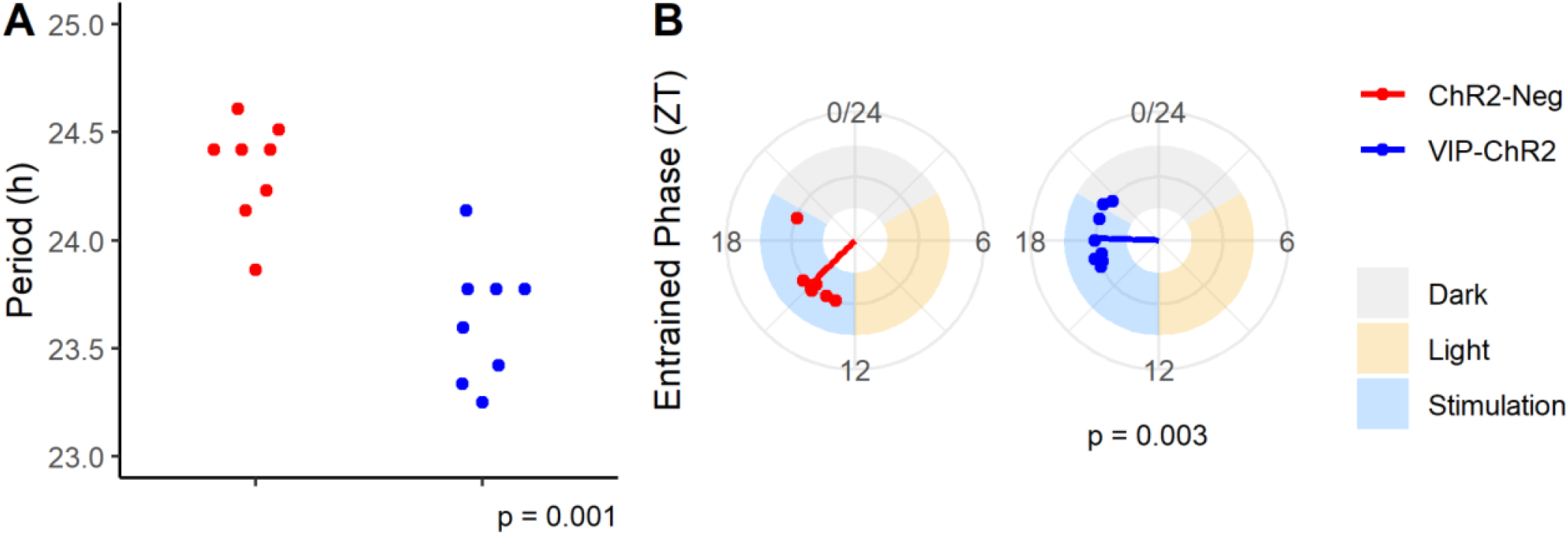
Optogenetic stimulation of VIPergic neurons influences *ex vivo* free-running period and entrained phase. A, free-running period of the PER2::LUC rhythm as measured by LSP on ~6 cycles of luminescence. The indicated *p* value is the result of a Kruskal-Wallis test. B, circular plots showing the first peak time of each slice in ZT. The indicated *p* value is the result of a circular ANOVA. Shaded regions represent the lighting conditions of the previous cycle at that phase.

To determine the phase angle of entrainment (the relative alignment of the SCN PER2::LUC rhythm with the entraining LD cycle), we determined the mean peak time of the first peak relative to the last lights-off transition before slicing (Figure 2B). The peak times for the ChR2-Neg group clustered in the hours following projected lights-off, and the peak times for the VIP-ChR2 group clustered in the second half of the projected stimulation window. The mean phase of the first peak in VIP-ChR2 slices (ZT 18.12, circular SD 0.42 h) was significantly delayed (*p* = 0.003, circular ANOVA) compared to that of the ChR2-Neg slices (ZT 15.02, circular SD 0.45 h), reflecting the phase delay induced by the repeated stimulation.

We performed a parallel experiment to determine if the plasticity we observed in the SCN *ex vivo* was also observable in behavioral changes *in vivo*. We measured locomotor activity of VIP-ChR2 and ChR2-Neg mice via passive infrared detection (PID). We exposed the mice to 5 days of short photoperiod (8:16 light:dark, h) followed by 7 days of 8 h light, 8 h stimulation in the dark, and 8 h of unstimulated darkness. We then untethered the mice and transferred the cages into constant darkness (DD) for 7 days (Figure 3A).

**Figure 3.**
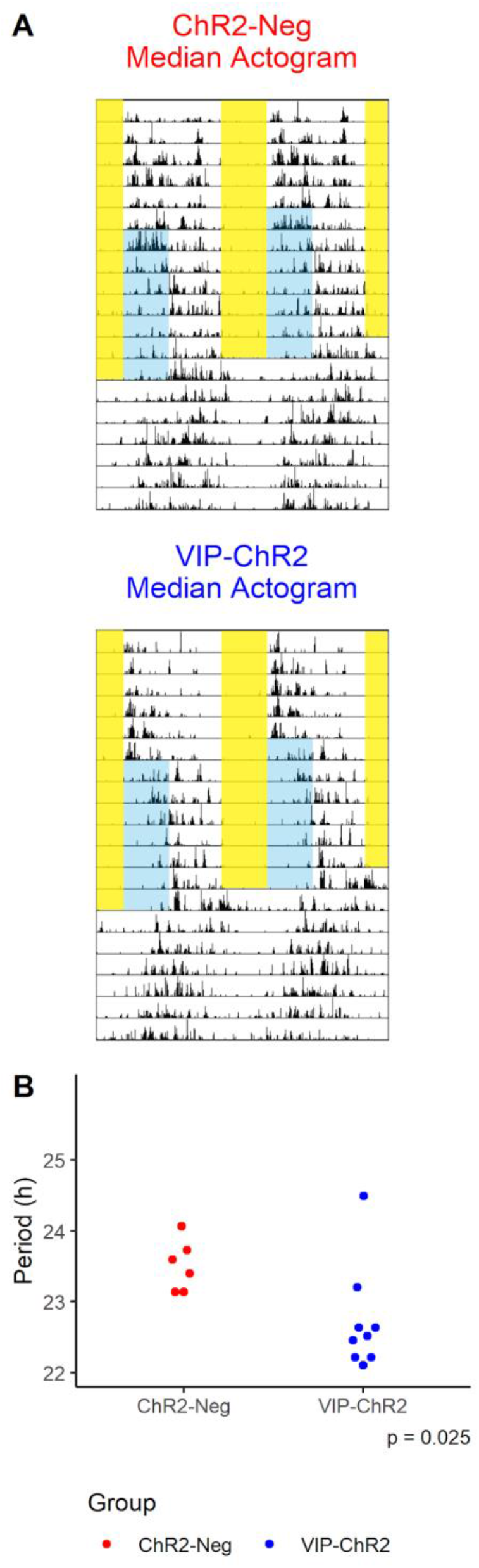
Optogenetic stimulation of VIPergic neurons influences *in vivo* free-running period. A, median actograms for ChR2-Neg (top) and VIP-ChR2 (bottom) mice. Yellow shading indicates light exposure, blue shading indicates blue light stimulation of the SCN (see also Figure S1). B, free-running period in each group as measured by LSP on cycles 2-7 of DD. The indicated *p* value is the result of a Kruskal-Wallis test.

Using the LSP on the DD portion of the activity record for each mouse, we found free-running period was significantly shorter in VIP-ChR2 mice (median 22.71, IQR 0.36 h) than in ChR2-Neg mice (median 23.72, IQR 0.25 h; *p* = 0.013, Kruskal-Wallis test; Figure 3B). This difference in period is similar in magnitude to that reported between long and short photoperiods [7,9] and the difference we observed *ex vivo* (Figure 2A), indicating that extended stimulation of VIPergic SCN neurons induces long-photoperiod-like free-running period lengths in mice housed in a short photoperiod both *in vivo* and *ex vivo*.

We also measured the effect of stimulation *in vivo* on phase angle of entrainment of the locomotor rhythm. Because the process of untethering mice and transferring the cages upon initiation of DD produced activity artifacts (Figure S3), we measured the phase angle using the full DD portion of the experiment as well as with the first 24 h of DD excluded. To estimate phase angle, we used a linear model to identify the peak of locomotor activity (“acrophase”) for each DD cycle, then estimated the acrophase of the last stimulation cycle by linear regression. This retrojected phase estimates the phase of the rhythm during entrainment without the direct influence of the light cycle and/or stimulation. When calculated excluding the first 24 h of DD, the mean phase of the VIP-ChR2 group was modestly more delayed (ZT 0.04, circular SD 1.09 h) compared to ChR2-Neg (ZT 21.13, circular SD 0.78; *p* = 0.186, circular ANOVA; Figure S4A), while phase estimates based on all 7 days of DD showed a large variance (Figure S4B). Differences in free-running period between the two groups were similar whether calculated using all 7 days of DD (Figure 2A) or excluding the first 24 h (Figure S4C).

The noisiness of the PID prevented reliable onset and offset detection, thus prohibiting us from conclusively quantifying a locomotor behavior duration (alpha) after-effect in either group. We observed, via qualitative analysis of the median counts in baseline and in stimulation, that mice in the VIP-ChR2 group, but not in the ChR2-Neg group, generally decreased their activity during the stimulation hours and increased their activity further into the dark phase upon the onset of the stimulation cycles of the experiment (Figure S5A, C). We were unable to determine if this pattern is maintained into the first cycle of DD, as the movement of the cages into DD produced activity artifacts (Figure S5E, top). The patterns are less noisy in the second cycle of DD, where we observed a median activity waveform approximately 2 h narrower at half-maximum in the VIP-ChR2 group compared to the ChR2-Neg group.

## Discussion

Our results elucidate the role of VIP in SCN photoperiodic encoding [4] and in the SCN light response [3]. We tested whether targeted activation of SCN VIPergic neurons can invoke persistent changes in SCN network configuration and function that constitute encoding of photoperiod in the isolated *ex vivo* SCN. Indeed, we found that optogenetic stimulation in the hours following lights-off in short, winter-like photoperiods induced changes in the SCN network similar to a true lengthening of the light cycle – re-aligning the phase of SCN molecular rhythms, increasing the phase dispersion of molecular rhythms across the SCN, and shortening the free-running period of the SCN clock [6,10–12].

Knockout of VIP disrupts the induction circadian responses to long photoperiod, the persistent compression of locomotor activity duration and the broadening of the SCN electrical activity waveform [4]. As such, VIP is thought to be necessary for the encoding and storage of photoperiodic information in the SCN. Here, we have shown that targeted daily optogenetic stimulation of VIPergic neurons timed to mimic long photoperiods can bring about the persistent increase in SCN network phase dispersion that underlies the broadening of electrical activity waveform and the compression of activity duration. The temporal precision of our stimulation demonstrates a specific role for VIPergic neurons in establishing the SCN network representation of photoperiod beyond previous experiments that have focused on the presence or absence of VIP itself.

In addition, we found that activation of VIPergic neurons was sufficient to establish additional persistent after-effects of long photoperiods in the SCN network, including a shortening of free-running period and altered phase alignment to dusk. Our lab and others have found that long and short photoperiods of 24 h T cycles have corresponding period after-effects *in vivo* and *ex vivo*, though the long-short difference is often small *ex vivo* [7,14,15]. Our results here, however, show a sizable decrease in period length for the VIP-ChR2 group (Figure 2A). The larger effect size suggests that extended stimulation of the VIPergic neurons may be a powerful influence on the free-running period of the SCN network, notably one that is more fully maintained from intact brain into slice culture. The shortened free-running period may itself be a consequence of the increased phase dispersion in the SCN [12,13]. Interestingly, VIP also appears to play a role in returning the SCN network to baseline from reconfiguration from photoperiod [14]. Taken together, our results and those of others suggest that VIP and VIPergic neuron activity are necessary and sufficient for the encoding and storage of photoperiodic information in the SCN network.

In our behavioral experiments, targeted stimulation of VIPergic neurons elicited the subsequent changes in phase and period expected from long photoperiods, but we did not observe changes in activity duration. This limitation may be due to the resolution of the method we used to measure locomotor behavior combined with the practical aspects of chronic optogenetic stimulation. Because our mice were tethered with an optogenetic fiber throughout behavior measurement, traditional running wheel activity measurement (with a cross-cage axle that would tangle the fiber optic tether) was not possible. We instead used passive infrared detection (PID), which has considerably lower signal to noise ratio. The resulting activity data was therefore difficult to analyze, and high-confidence manual scoring of activity duration (alpha) was not possible. In the future, it may be possible to use axle-free running wheels that provide the high signal-to-noise ratio of traditional running-wheel actigraphy without the risk of interference with the fiber optic tether to determine the alpha after-effects following extended VIPergic SCN neuron activation.

Some optogenetic studies use enucleated mice to avoid retinal activation from fiber optic light leak[5]. Our experiment, which required light stimulation in addition to optogenetic stimulation, precluded enucleation. We used cladded fiber optic tethers, but fiber junctions at the base of the fiber optic implant and between the implant and the tether allow some light leak. We controlled for this issue by stimulating both our experimental (VIP-ChR2) and control (ChR2-Neg) mice identically, so that any effects of retinal activation on circadian behavior would be visible in the control group. In fact, some negative masking may have occurred in some control animals during the stimulation window (Figure 3A, top; Figure S3), but any effects disappear in DD (hence “masking”) in all but one control mouse (OC012, Figure S3). In experimental mice (Figure 3A, bottom), the negative effect on activity during the stimulation window appears stronger and persists into DD in many of the mice. The groups responded differently to the treatment despite identical surgery and stimulation procedures as shown by their differences in phase following treatment *in vivo* (Figure S4A) *ex vivo* (Figure 2B). If retinal stimulation was responsible for our observations, we would have likely seen similar phase in both groups

By demonstrating the sufficiency of VIPergic neurons in the induction of circadian photoperiodic encoding, we have further established the importance of this neuronal subpopulation in transducing photic signals from the retina to the rest of the SCN and downstream to the rest of the body. If stimulating these neurons is sufficient to induce photoperiodic effects, can inhibiting the neurons block the effects? Optogenetic inhibitory tools are currently not as effective as those that induce neural activity, but chemogenetic approaches may be better suited to answer this question in the future.

Here we focused on circadian photoperiodic changes, which are reliable, clock-related changes that can be observed even in mice that, like many common lab strains, lack appreciable melatonin synthesis. We have previously shown that photoperiod exposure in melatonin-competent mice can induce more sophisticated changes in physiology, such as depression-anxiety behavior [15]. In the future, it will be helpful to determine whether the sufficiency of VIPergic neurons in inducing circadian photoperiod after-effects translates to the induction of such additional down-stream effects, as it would provide a specific neuronal target for possible therapeutic manipulation.

## Acknowledgments

We would like to thank Jeff Jones and Elliot Outland for helpful discussions. This work was supported by NIH R01 GM117650 to D.G.M. and NIH R35 GM124685 to J.J.H.

## Author Contributions

M.C.T. and D.G.M. designed the experiments. M.C.T. performed the experiments. M.C.T. and J.J.H. analyzed the data. M.C.T., J.J.H., and D.G.M. wrote the paper.

## Declaration of Interests

The authors declare no conflicts of interest.

## STAR Methods

### Resource Availability

#### Lead contact

Further information and requests for resources and reagents should be directed to, and will be fulfilled by, the lead contact, Douglas G. McMahon

#### Materials availability

This study did not generate new unique reagents.

#### Data and code availability

The code generated during this study is available at FigShare: https://doi.org/10.6084/m9.figshare.13519043

### Experimental Model and Subject Details

#### Mouse Lines

For *in vivo* experiments, 5 *Vip::Cre*^+/−^*Per2::Luciferase*^+/−^ (4 male, 1 female) and 1 *Vip::Cre*^+/−^ (male) mice were used as controls (ChR2-Neg). 9 *Vip::Cre*^+/−^*floxed-ChR2*^+/−^ (7 male, 2 female) were used as the experimental group (VIP-ChR2).

For *ex vivo* experiments, 8 *Vip::Cre*^±^*Per2::Luciferase*^+/−^ (5 male, 3 female) mice were used as controls (ChR2-Neg). 8 *Vip::Cre*^+/−^*floxed-ChR2*^+/−^*Per2::Luciferase*^+/−^ (4 male, 4 female) mice were used as the experimental group (VIP-ChR2).

### Method Details

All procedures involving mice were performed in accordance with Vanderbilt University Institution of Animal Care and Use committee regulations.

#### Fiber optic implant surgery[16]

We anesthetized mice with 3% isoflurane and provided ketoprofen for pre- and post-operative analgesia. After isoflurane induction, we secured the head of the mouse in a streotax and provided lubricating eye drops. We applied iodine-based antiseptic to the shaved scalp and made an incision in the skin. We applied 2% hydrogen peroxide to clean the skull of connective tissue and dried the surface with a sterile cotton swab. After leveling the skull, we used a mounted drill to make a small craniotomy at Bregma and scoured the skull surface with forceps. We lowered a 5 mm fiber optic implant into the craniotomy and stopped any bleeding that results with ophthalmic absorbent strips. We then applied Metabond to the scoured skull surface surrounding the fiber optic implant post and let it cure for 5 minutes, followed by a small but secure dental cement cap that was allowed to cure for 10 additional minutes. We secured the skin to the dental cement cap using sterile veterinary surgical adhesive and coated the edges of the skin with iodine-based antiseptic.

#### *Ex vivo* procedure

After at least 1 week of recovery from surgery in 12:12 LD, we moved the mice to the stimulation light/dark box, still in standard shoebox cages (no running wheel, no PID attachment) so that no acclimation was necessary before tethering. We immediately attached these mice to the fiber optic tethers and stimulation began at the time of lights-off that evening. After 7 days of daily stimulation, we extracted and cultured 300 μm slices of SCN for PER2 luminescence recording.

We made SCN slices 2-4 h before the timing of lights-off. For slicing, we used cold, HBSS-based media containing antibiotics, HEPES, and sodium bicarbonate. Our slice culture media contained 0.1 mM of luciferin, D-glucose, HEPES, antibiotics, sodium bicarbonate, and standard B-27. We cultured the slices in a six-well plate with 1.2 mL of culture media in each well, with a sterile culture membrane on top. We sealed the plates with PCR plate sealing film and imaged the slices using an inverted microscope with an intensified CCD camera attached. We imaged each slice once per 10 minutes, using an aggregation time of 2 minutes, and continued recordings for 6 days.

#### *In vivo* procedure

We used a separate cohort of mice undergoing the same surgical and recovery procedures for our *in vivo* studies. After recovery in 12:12 LD, we moved animals to 8:16 LD (short photoperiod) in cages with passive infrared detectors (PIDs) attached. There were no running wheels in the recovery or experimental cages. We housed mice for 2 days without attachment to the fiber optic tethers to acclimate to the PID cages. On the second full day, we attached mice to the fiber optic tether and the following 5 days were designated as the “Baseline” portion of the experiment. On the 6th day, the LEDs began turning on for 8 h each day at 10 Hz, starting at lights-off. To avoid any effect of a dark pulse between the termination of light and the onset of stimulation, we set the stimulation to overlap the light cycle by ~ 3 minutes. After 7 full days of daily stimulation, we moved the cages of the mice into a nearby light/dark box for the constant darkness (DD) portion of the experiment. Cage transfers were done in dim red light. After 7 full days in DD, we ended the experiment.

Two mice (one each from ChR2-Neg and VIP-ChR2) were re-run through the *in vivo* procedure due to an LED failure and a PID failure, respectively. Neither of these mice were transferred to DD, but rather remained in the stimulation light/dark box for the following run-through of the protocol. Neither mouse received fiber optic stimulation during their initial, failed trial, but did receive extra days of 8:16 photoperiod as a result.

### Quantification and Statistical Analysis

#### *Ex vivo* analyses

Each SCN slice image corresponded to ~6 cycles of PER2::LUC bioluminescence with a resolution of 6 frames per h. We processed each image in ImageJ first using a two-frame minimization to remove CCD noise (reducing frame rate to 3 frames per h) followed by an overlay of a grid of 10,404 ~10 × 10 μm ROIs. We exported the “stack grid” of the luminescence profiles of each ROI into R for further analysis. We limited our analyses to ROIs within the top 20th percentile of overall luminescence intensity in each slice, thus focusing the analyses on the SCN tissue. For the PER2::LUC trace in each ROI, we smoothed using a Savitzky-Golay filter (order of 2, span of 51) and baseline subtracted using a more heavily-smoothed Savitzky-Golay filtering of the trace (order of 5, span of half the number of frames). For each baseline-subtracted, smoothed trace, we identified up to 6 peaks (at least 18 h apart).

To eliminate artificial or partial first peaks, we established a starting cutoff at 18 h post-culture (Figure S4A). From that starting cutoff, we used a sliding 24-h window to assign peak numbers to each detected peak regardless of the order identified (this prevents a missed peak from altering all subsequent peak times, Figure S4B). For example, if ROI A identified peaks at 24, 48, and 72 h and ROI B identified peaks at 24 and 72 h, this step will appropriately designate ROI A’s peaks as Peak 1, Peak 2, and Peak 3 and ROI B’s peaks as Peak 1 and Peak 3.

Using this processing, we calculated the mean peak time of each overall peak and calculated the phase of each ROI relative to the mean timing of that peak for that slice. For each peak in each slice, we calculated the median absolute deviation of peak times and performed an ordinary two-way ANOVA to measure the main effect of peak number and group.

With the same starting cutoff of 18 h, we measured free-running period of the PER2::LUC rhythm using the Lomb-Scargle periodogram. We compared the groups using a Kruskal-Wallis test on ranks.

We measured phase as the mean time of the first peak for each slice. Like our *in vivo* analysis, we compared the phase of the two groups using a circular ANOVA, while calculating the circular mean and standard deviation of each group.

#### *In vivo* analyses

We generated median actograms using ClockLab Analysis (Actimetrics). We estimated the acrophase for each mouse in each cycle following the final stimulation offset. In R, we fit a linear regression of the acrophase for cycles 1-7 or cycles 2-7 to retroject the acrophase of the final day before DD. We analyzed phase differences between the two groups using a circular ANOVA with the circular package. Using the same package, we calculated circular means and standard deviation.

We measured free-running period using the Lomb-Scargle periodogram on the counts (1-minute resolution) for cycles 1-7 (Figure 3B) or cycles 2-7 (Figure S4C) from each actogram exported to R. We analyzed the free-running period using a Kruskal-Wallis test on ranks.

## Supplemental Information Titles/Legends

**Figure S1.**
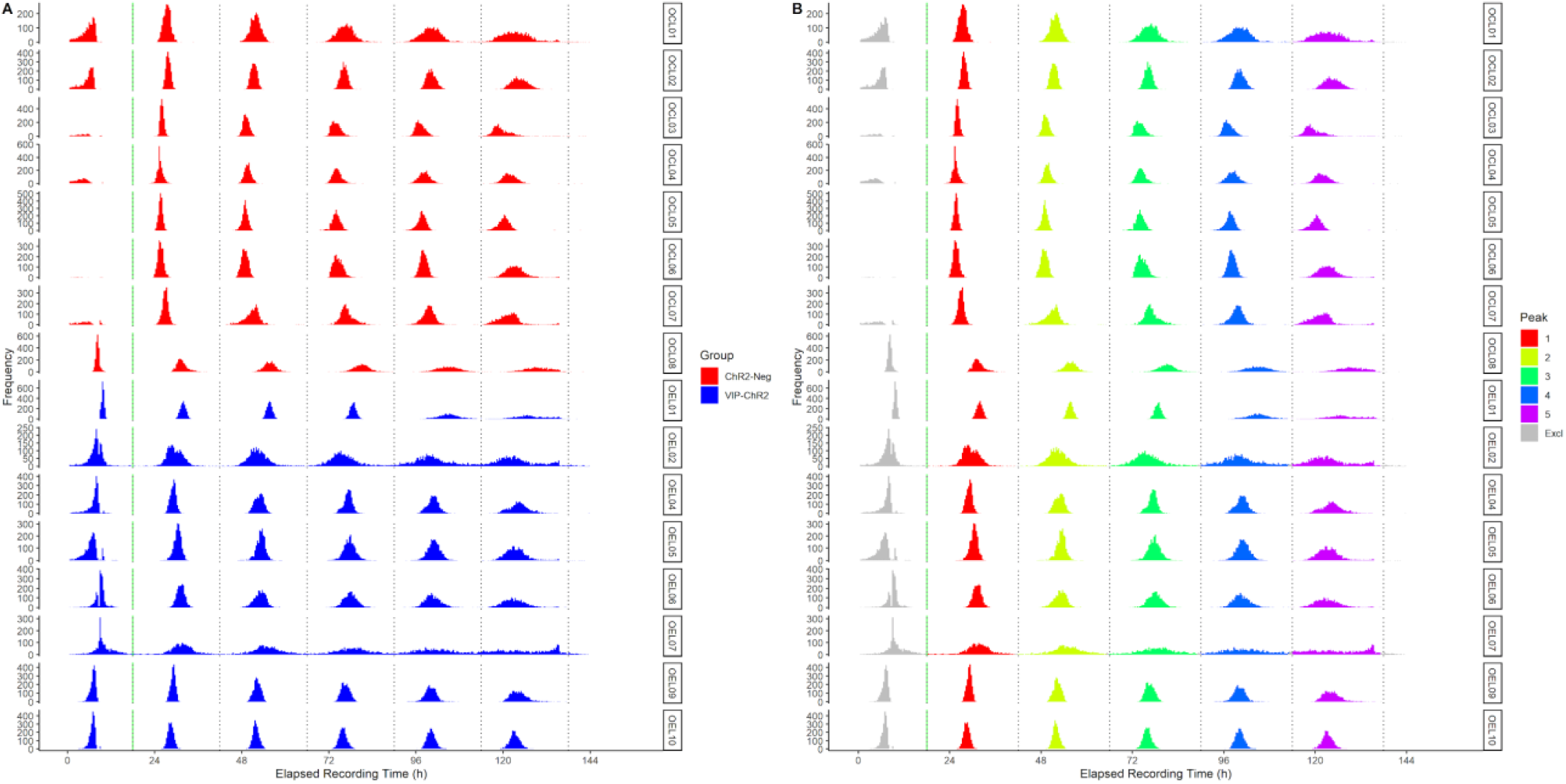
(related to Figure 1). Frequency distributions for the detected peak times for all SCN ROIs for each slice. A, raw peak times for each slice with ChR2-Neg slices in red and VIP-ChR2 slices in blue. Green line represents the starting cutoff for inclusion (see Methods). Subsequent black lines represent 24 h intervals beginning at the starting cutoff used for peak number assignment in B. B, the peak number assignments from the ROI peak times shown in A. Based on the 24 h intervals beginning with the starting cutoff (green line), peaks occurring within each successive interval were assigned to the specified peak number.

**Figure S2.**
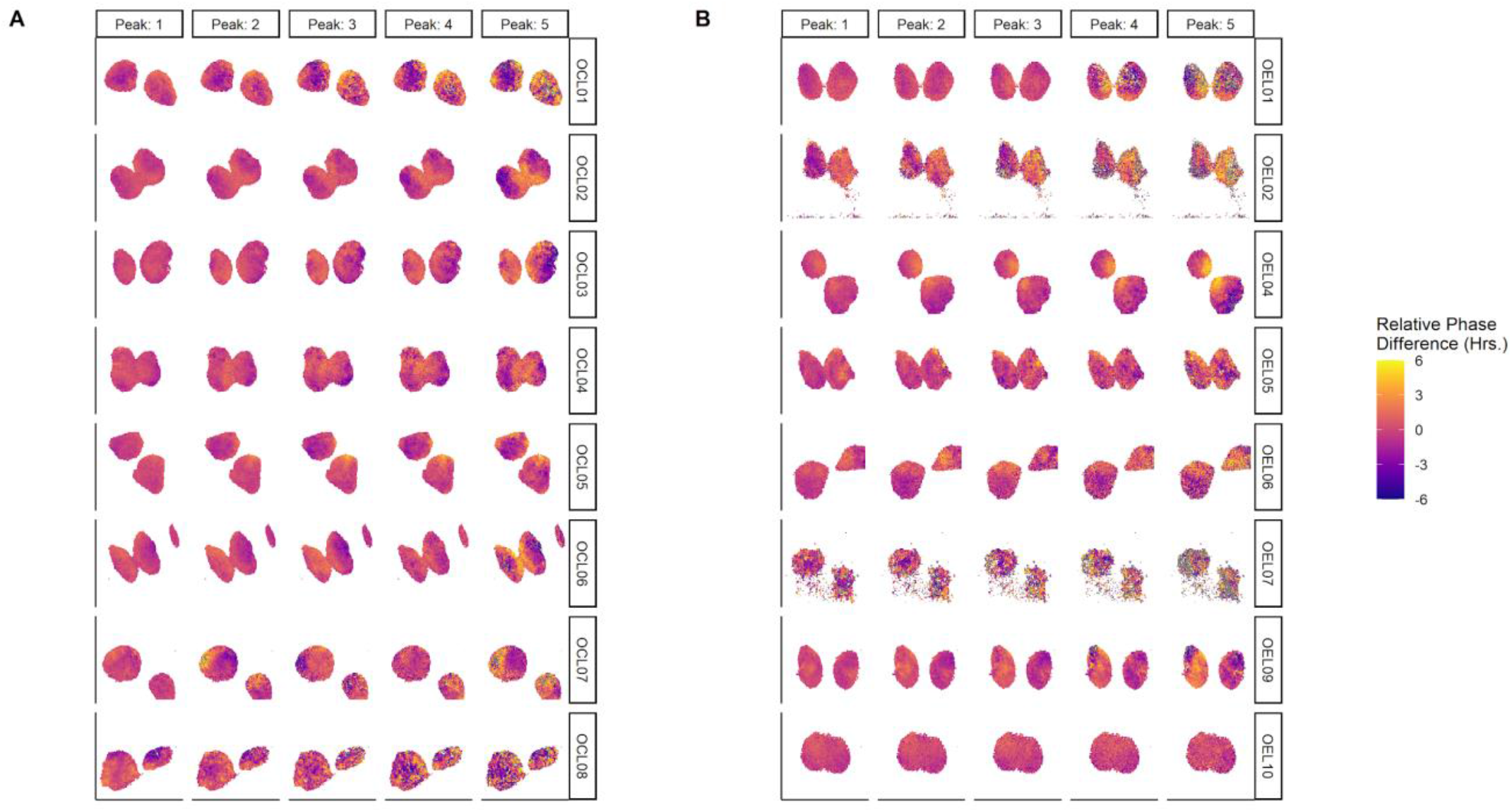
(related to Figure 1). Individual relative phase maps for each slice used in the *ex vivo* analyses. A, relative phase maps for the ChR2-Neg group. B, relative phase maps for the VIP-ChR2 group. In all maps, the relative phase is calculated as the difference, in hours of each ROI relative to the mean phase of the slice at that peak. Warmer colors represent delayed phase, cooler colors represent advanced phase.

**Figure S3.**
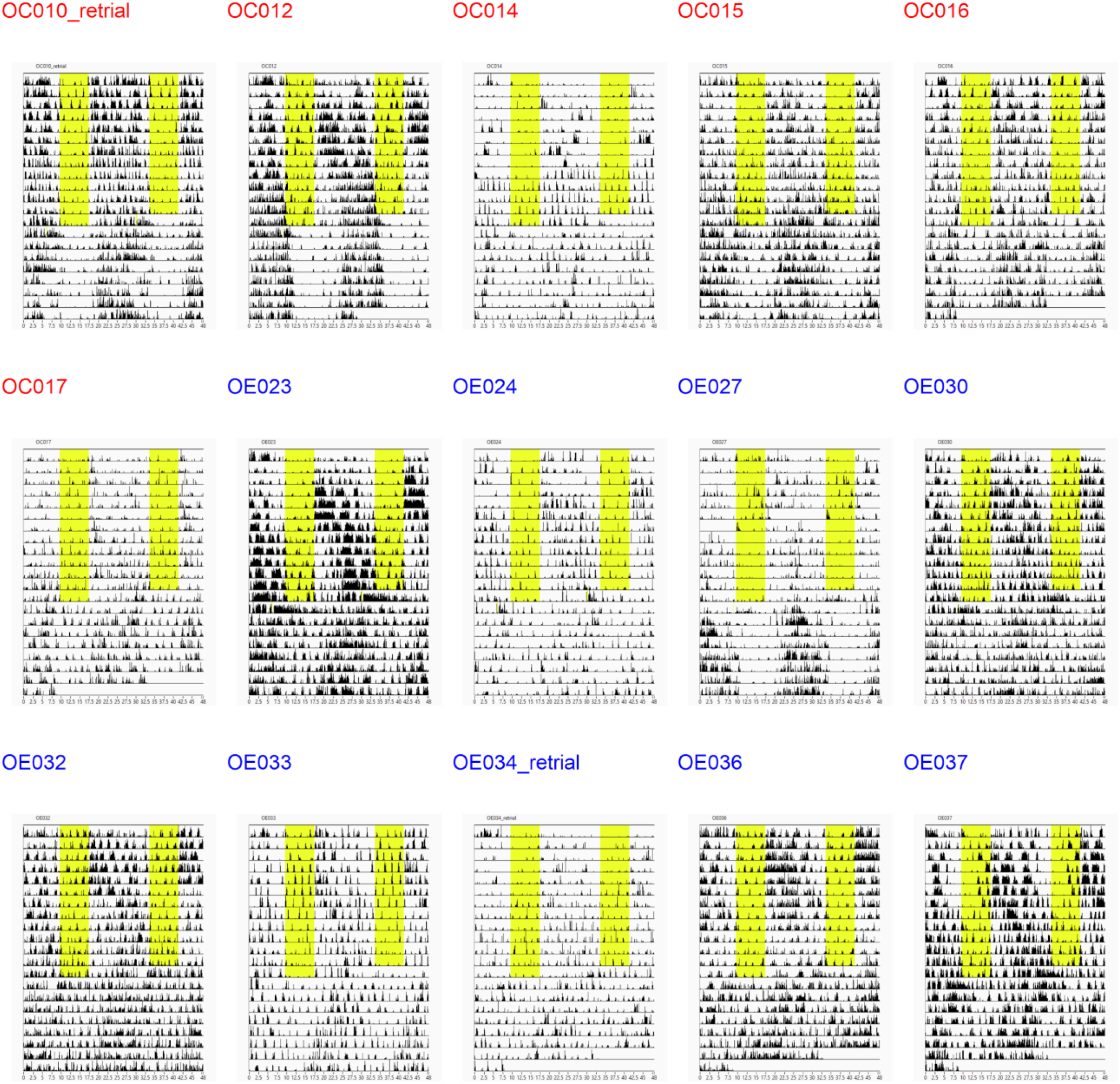
(related to Figure 3). Individual actograms for each mouse used in the *in vivo* analyses. Actograms are double-plotted with 6-minute binning. Yellow region represents 8 h window of daily light exposure. Stimulation window (not shown) was 8 h following lights off on the final 7 days of light exposure. Animals were attached to fiber optic implants on the second day of the records.

**Figure S4.**
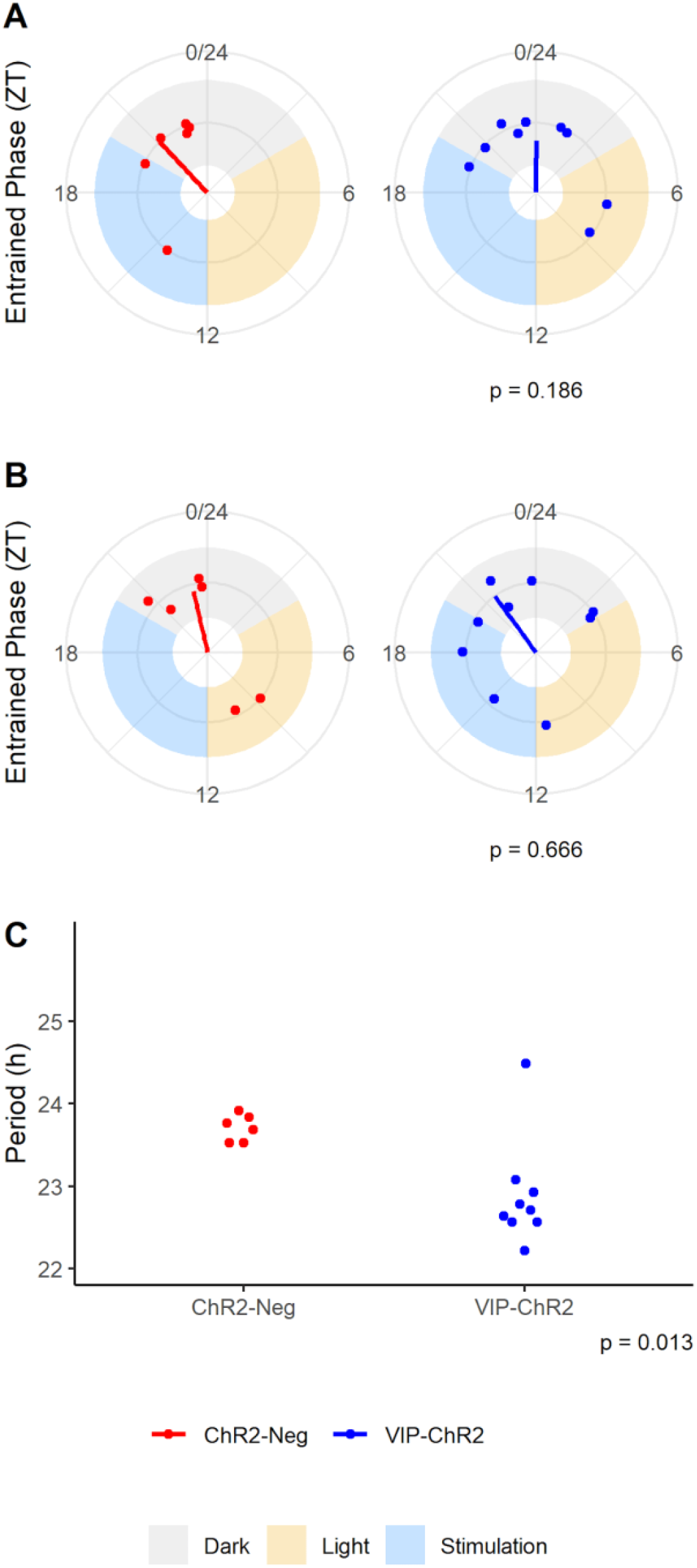
(related to Figure 3). Entrained phase of the locomotor behavior rhythm for each mouse estimated excluding the first 24 h of DD (A) and using all 7 days of DD (B). The free-running period calculated excluding the first 24 h of DD.

**Figure S5.**
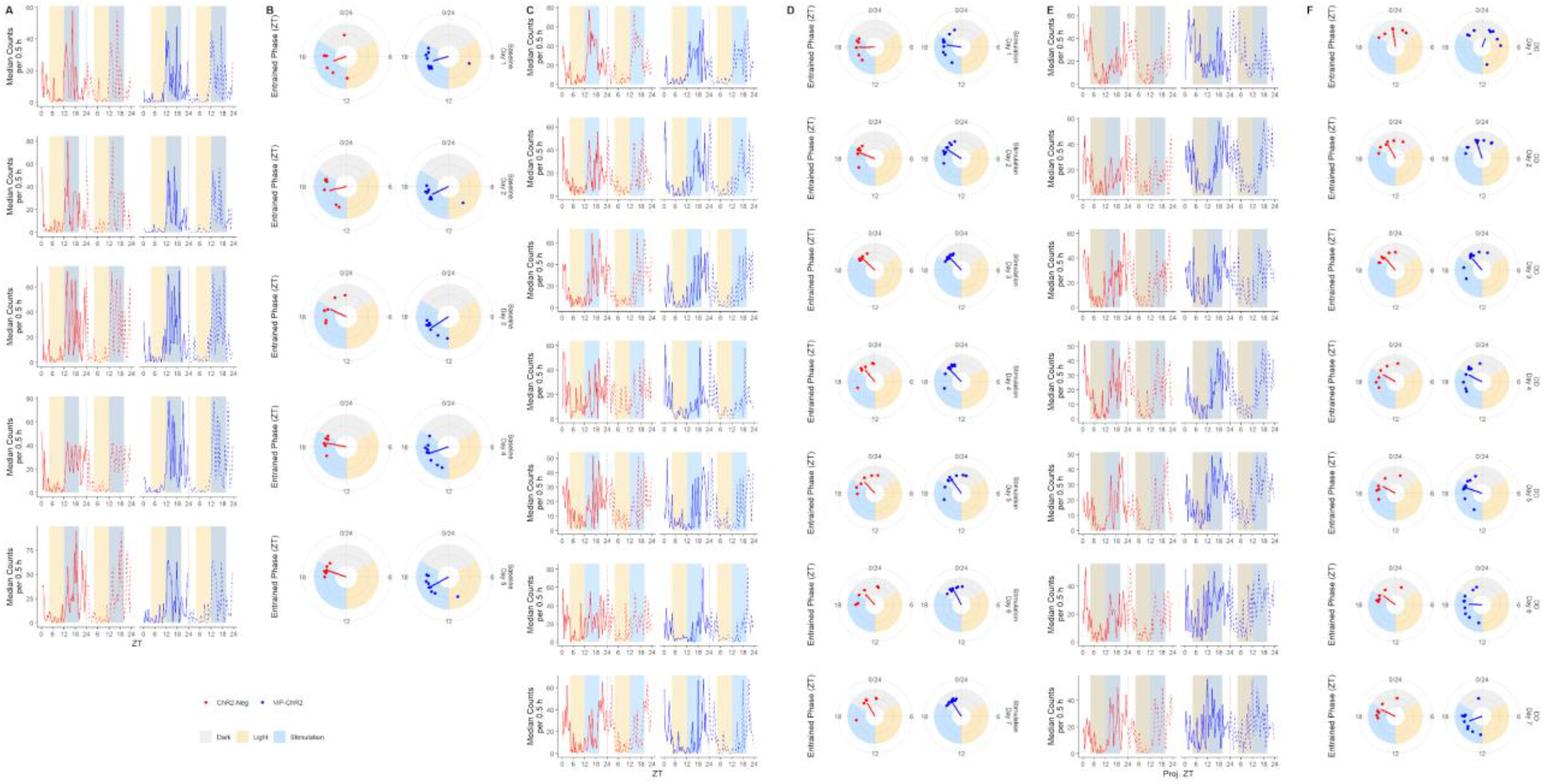
(related to Figure 3). Double-plotted median half-hourly-counts (A, C, E) and associated acrophase estimation (B, D, E) for each group in each day of Baseline (A, B), Stimulation (C, D), and DD (E, F). Yellow shading indicates light, blue shading indicates LED stimulation of the SCN. Dark blue represents the stimulation window on non-stimulation days (Baseline, DD), dark yellow represents the light window on DD days. Indicated *p* values are calculated by circular ANOVA. Dashed lines represent double-plotted data.

## Notes

### Competing Interest Statement

The authors have declared no competing interest.

